# A Systematic Protein Turnover Map for Decoding Protein Degradation

**DOI:** 10.1101/2020.03.09.983734

**Authors:** R. Christiano, S. Kabatnik, N. Mejhert, R.V. Farese, T.C. Walther

## Abstract

Protein degradation is mediated by an expansive and complex network of protein modification and degradation enzymes. Matching degradation enzymes with their targets and determining globally which proteins are degraded by the proteasome or lysosome/vacuole has been a major challenge. Further, an integrated view of protein degradation for cellular pathways has been lacking. Here we present a novel analytical platform that combines systematic gene deletions with quantitative measures of protein turnover to deconvolve protein degradation pathways for *S. cerevisiae*. The resulting turnover map (T-MAP) reveals target candidates of nearly all E2 and E3 ubiquitin ligases and identifies the primary degradation routes for most proteins. We further mined this T-MAP to identify new substrates of ER-associated Degradation (ERAD) involved in sterol biosynthesis and to uncover novel regulatory nodes for sphingolipid biosynthesis. The T-MAP approach should be broadly applicable to the study of other cellular processes and systems, including mammalian systems.

**One Sentence Summary:** A systematic, global map of protein turnover for a large set of yeast mutants reveals scope and specificity of degradation pathways and identifies novel regulatory nodes for lipid metabolism.

---

The abundance and composition of specific proteins are key determinants of cell function. Degradation is crucial for controlling the abundance of many proteins, and it is mediated by a large network of protein modification, sorting, and degradation machineries. Principally, proteins are degraded by either the proteasome (*1*) or acid proteases in the lysosomal compartment (the vacuole in yeast) (*2, 3*). Covalent modification with the small protein-modifier ubiquitin (*4, 5*) through the sequential actions of E1, E2, and E3 ligases is important for selecting substrates for either pathway. However, many mysteries remain about protein degradation, such as the routes of degradation taken by each protein individually, the identity of ubiquitin ligases that modify specific proteins, and how protein degradation pathways are integrated to regulate cell function. A major hurdle has been developing strategies to measure protein turnover globally, in a fashion similar to strategies that exist for measuring cellular mRNA expression levels (by RNAseq or microarrays) or protein synthesis (by ribosome footprinting).

With recent progress in mass-spectrometry-based proteomics, unraveling these mysteries is now within reach. For example, global analyses of protein turnover in *Saccharomyces cerevisiae* and *Schizosaccharomyces pombe* revealed that proteins lifetime fall into three classes (*6, 7*). Most proteins in budding yeast (~85%) are long-lived (t_1/2_ >5 hrs), and their amounts are determined chiefly by their rates of synthesis and dilution. The remaining short- (t_1/2_ <1.25 hrs) and medium-lived proteins (1.25 hrs< t_1/2_<5 hrs) are actively degraded at steady-state, controlling their abundance. Turnover of this ~15% of the proteome is particularly important as this class is rich in regulated factors such as cell cycle proteins, membrane transporters or metabolic enzymes (*6*). The ability to globally monitor turnover of native proteins (endogenous promoters and no tags) with minimal cellular perturbations (no drugs) opens the door to performing systematic turnover profiling studies in combination with genome perturbations to solve many longstanding questions pertaining to protein degradation. In particular, it is now possible to combine monitoring protein turnover rates across series of genetic deletions to elucidate biochemical networks.

To comprehensively understand protein degradation, we developed an approach to measure protein turnover in a large set of *S. cerevisiae* mutants deficient in specific components of the protein degradation machinery. We utilized state-of-the art proteomics, combined with pulse-labeling of yeast with amino acids containing stable isotopes, to globally measure endogenous protein turnover in yeast strains with mutations of ~120 genes encoding components of the protein degradation machinery (**Fig. 1A and Suppl. Fig. S1A**). Our previous study showed 60 min after pulse-labeling to be the optimal single time for turnover analysis of most proteins (*6*), and we used this timepoint for our measurements. This strategy accurately captured changes for most proteins in a resultant turnover map (T-MAP), comprising 465,510 protein abundance and turnover measurements from 4,585 proteins (derived from 86,502 peptides identified at 1% FDR). To ensure the accuracy and reproducibility of our measurements, we implemented a series of quality control strategies, such as independent biological duplicate measurements and testing for absence of mass spectrometry signals from peptides encoded by the deleted gene (see supplementary information; **Fig. S1C and S1D**). Analyzing the median of all measurements across mutants for each protein as a proxy of their turnover in a wildtype setting (see supplementary information; **Fig. S1C**) revealed that each organelle contains short-, medium- and long-lived proteins, suggesting that protein turnover is determined for proteins individually, rather than for a whole organelle (**Fig. S1E**).

**Figure 1:**
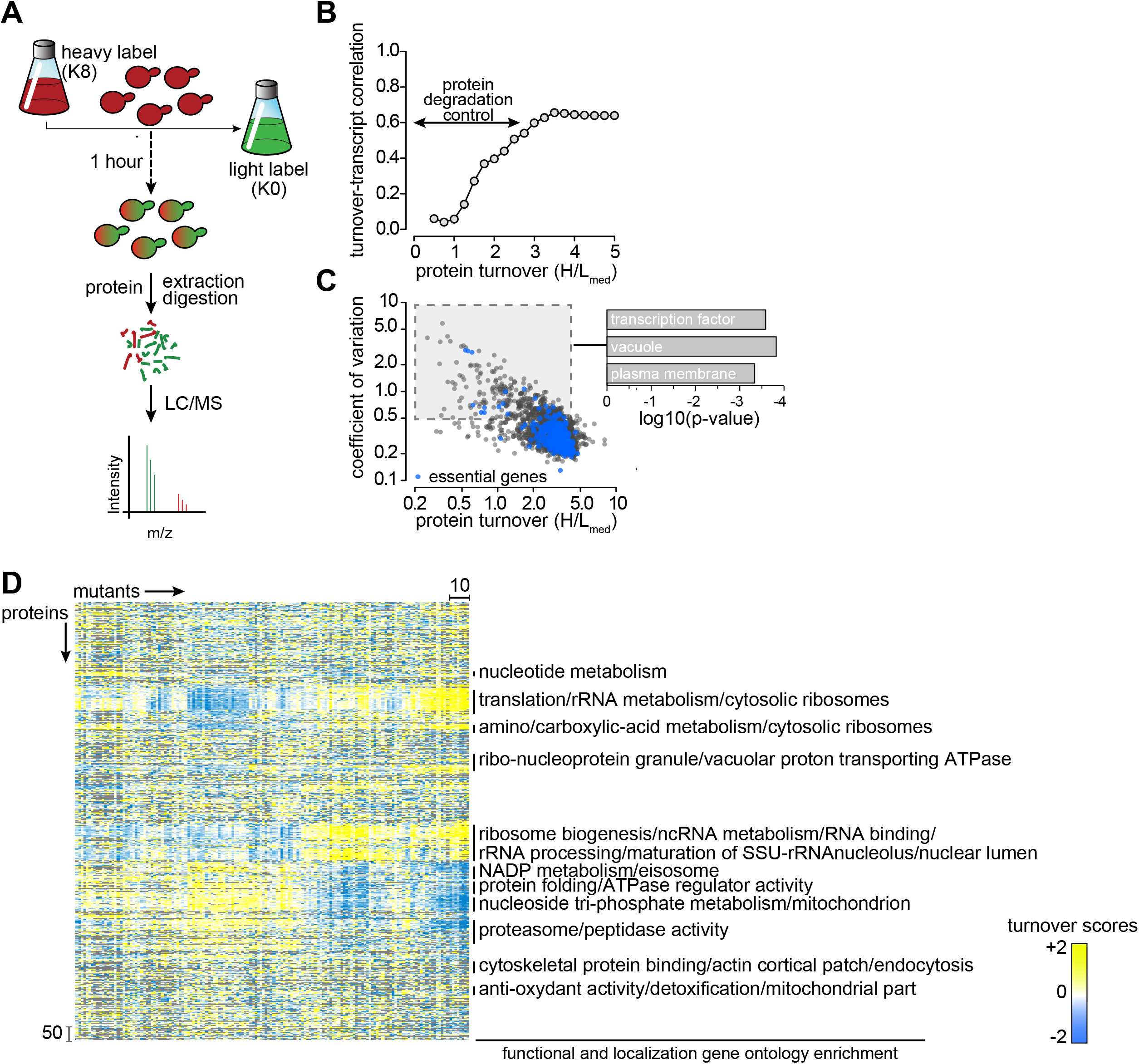
Overview of the Turnover Profiling Map (T-MAP) of Proteostasis Genes. **(A)** Overview of single time point strategy for turnover profiling. **(B)** Distribution of Pearson correlation coefficients between pairwise turnover correlation and expression correlation (transcript abundance; (Kemmeren et al., 2014) as a function of protein turnover. **(C)** Coefficients of variation of H/L ratios of each protein plotted as a function of the median of H/L ratios (H/Lmed). Essential proteins are in blue; other proteins are in black. **(D)** Hierarchical clustering of proteostasis mutants (left-right) and proteins (top-bottom). Enrichment for functional and localization categories in clusters is indicated on the right.

Whether changes in mRNA levels accurately predict changes in the proteome has been a long-standing question (*8*). Integrating turnover data with measurements of mRNA transcript changes in a subset of our mutants (*9*) showed that, for long-lived proteins with slow degradation rates, transcriptional regulation accurately predicted protein turnover and abundance changes (**Fig. 1B)**. However, transcriptional changes in response to genome perturbation did not predict turnover and protein amount for short-lived and medium-lived proteins. Instead, degradation had a major effect for these classes of proteins. This group of short- and medium-lived proteins contained proteins with the highest turnover variation due to genome perturbation (c.v.>=50%) and was enriched for transcription factors (p-value=2.6 x10^−4^), plasma membrane proteins (p-value=1.5 x 10^−4^), and vacuolar proteins (p-value=4.6 x10^−4^), likely reflecting the importance of these proteins for maintaining cell homeostasis (**Fig. 1C)**.

To determine the effects of genetic perturbations on protein turnover, we developed a scoring function (robust T-score) to measure stabilization and destabilization effects (**Fig. S1C)**. An overview of the T-MAP scores in hierarchical clusters of protein turnover profiles, detecting classes of co-regulated proteins, is shown in **Fig. 1D** (see also **TreeView, Supplementary Tree1**). This analysis shows a coordinated response in the proteome turnover of major protein synthesis pathways (e.g., rRNA processing, ribosome biogenesis, and translation) and anabolic pathways (e.g., NADP metabolism and nucleotide metabolism). In addition to sorting co-regulated proteins, the T-MAP provides comprehensive information on the turnover profiles of nearly all yeast proteins for each deletion mutant (**Fig. 1D** and **TreeView, Supplementary Tree1**).

An important, open question for global protein turnover is which proteins are degraded by vacuolar versus proteasomal degradation pathways. We addressed this question by determining the effects of mutations in *PEP4*, the major vacuolar protease, and *RPN4*, a transcription factor that drives expression of most proteasomal subunits (*10, 11*), on the turnover of endogenous proteins **(Fig. S2A and S2B**). For the proteasomal degradation pathway, analyses of proteins stabilized in *rpn4D* cells revealed 128 short- and medium-lived proteins (**Table S4**) that are enriched in cell cycle–related proteins, kinases, ribosomal proteins, ubiquitination machinery, and some ER proteins (**Fig. 2A and 2B**). In contrast, many proteins of the secretory and endolysosomal compartments were enriched among the 106 short- and medium-lived proteins stabilized in *pep4D* cells, indicating degradation by the vacuole (**Table S4**; **Fig. 2A and S2C**). Unexpectedly, we also found soluble and membrane proteins from mitochondria that were stabilized in *pep4*D cells, suggesting continuous degradation of specific mitochondrial parts in the vacuole. Recently, a pathway was reported that removes specific mitochondrial membrane proteins in aged cells (*12*); our data now suggest a similar pathway may operate in rapidly growing yeast cells when mitophagy does not occur. We also identified 14 proteins that were stabilized in both vacuolar and proteasomal degradation mutants **(Fig. S2F)**, likely reflecting separate pathways for degrading distinct pools of protein. For example, Chs3 is present on the vacuolar membrane where it is partially internalized and degraded (*13*) but also on the ER from where it might be delivered to the proteasome.

**Figure 2:**
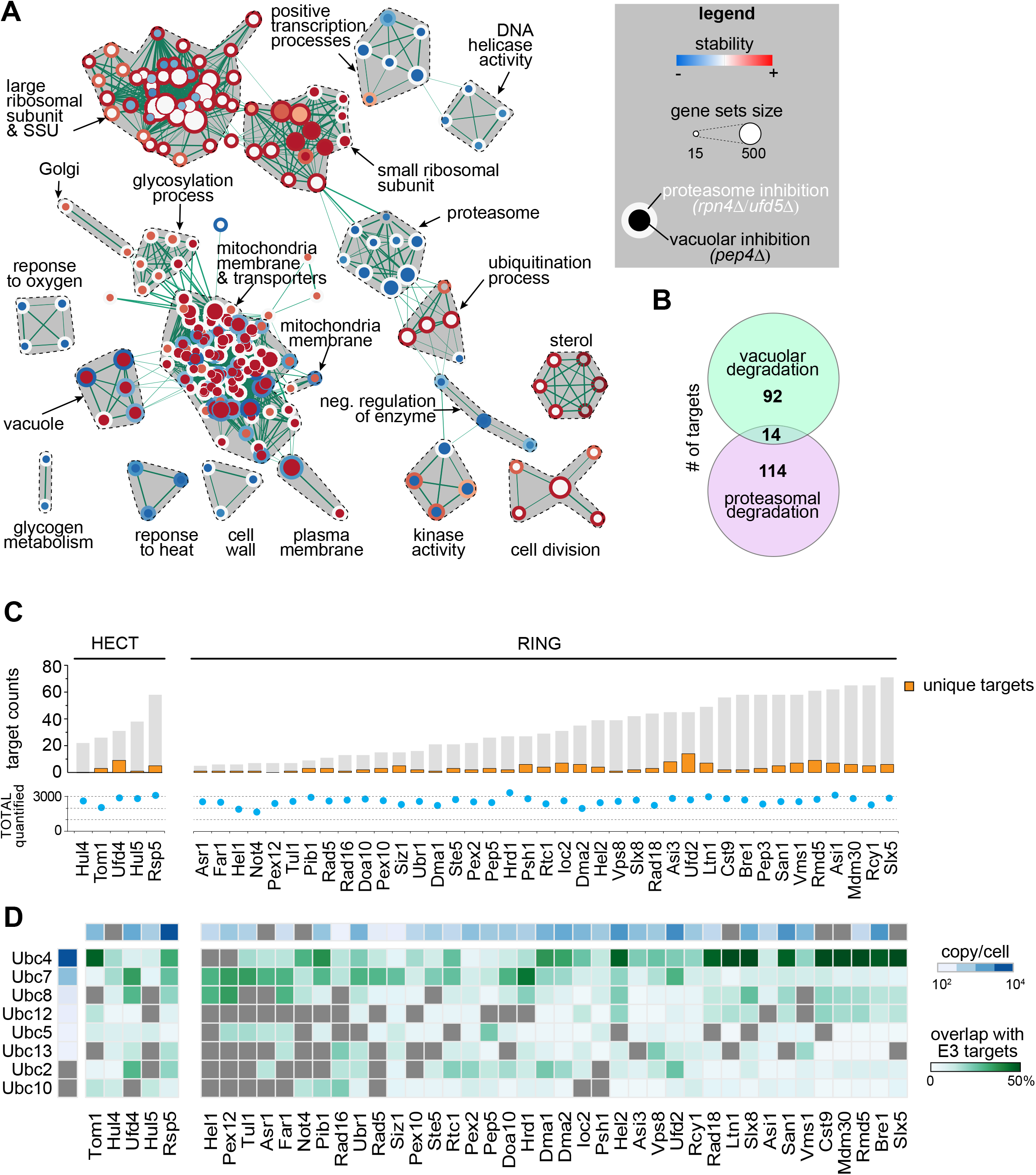
Turnover Effects of Lysosomal and Proteasomal Pathway Perturbations. **(A)** Enrichment map of Gene Set Enrichment Analysis (GSEA) for proteins affected by inhibition of proteasomal and lysosomal degradation (p=0.005, q=0.1). Nodes represent gene sets that are enriched at the top or bottom of the ranking of differentially affected proteins (as determined by GSEA). Node size corresponds to the number of genes in the set. Edges indicate overlap between gene sets, and the thickness indicates the size of the overlap. Red indicates increased and blue indicates decreased turnover. Node center corresponds to effect of vacuolar degradation inhibition (*pep4*Δ cells), whereas node rims correspond to the effects of proteasomal degradation inhibition (*rpn4*Δ cells). **(B)** Venn diagram representing the number of vacuolar and proteasomal targets. **(C)** Number of targets of HECT and RING E3 ligases present in the T-MAP. Unique targets are indicated in orange. Shared targets are indicated in grey (shared between at least two E3 ligases). **(D)** Target overlap between E3 and E2 enzyme s (in % of E3 ligases total targets, green). E3 and E2 enzyme abundances are indicated (blue).

Our analyses unexpectedly also revealed coordination of the proteasomal and vacuolar protein degradation pathways. For instance, all proteasomal subunits were turned over faster in *pep4*Δ cells (**Fig. 2A and S2D**). Conversely, many vacuolar enzymes (e.g., Pep4, CPY/Prc1, Prb1, Cps1, Ape3 and Pho8) turned over faster in *rpn4Δ* cells (**Fig. S2E**). Our understanding of the reasons and mechanisms for such coordination remains elusive but such questions can now be addressed in further studies with more genetic perturbations.

A major goal for the field has been to identify the targets for specific ubiquitin ligases. Towards this goal, we systematically mutated the entire set of non-essential E3 and E2 ligases, expecting short-lived substrates to be stabilized in the absence of their cognate E2 and E3 enzymes. Consistently, we found the vast majority of short- and medium-lived proteins are stabilized in at least one E3 ligase deletion strain. Investigating specific cases, we found our data to recapitulate prior findings on the requirements of substrates for particular E3 ligases, thus validating our dataset (**Suppl. Fig S2G**). In addition, we identified new targets for many E3 enzymes, with many proteins that apparently access multiple E3 ligases, including 96 proteins (21%) that are targeted by five or more E3 ligases **(Fig. S2H).** Conversely, mutations of some E3 ligases stabilize only few targets (e.g., Hel1 has six identified targets), but other E3s target as many as ~70 proteins (e.g., Slx5) **(Fig. 2C)**. Cooperation between E2 and E3 enzymes appears to be loose, inasmuch as there is great overlap between any E2 and multiple E3 targets (**Fig. 2D**). The most abundant E2 enzyme Ubc4 has more overlap with E3 enzymes than less abundant E2 enzymes, such as Ubc10 or Ubc13, suggesting that cooperation between E2 and E3 enzymes might be driven to some extend by mass action instead of specific protein-protein interactions (**Fig. 2D)**. Beyond these insights, our analyses now delineate the specificity for interactions in the ubiquitin proteasome system, allowing for further mechanistic dissection of the network and for defining the rules that govern access to specific E3 ligases using the new targets we have now identified.

For instance, previous work showed that the Hrd1 E3 ligase complex (**Fig. 3A**) targets misfolded substrates that contain lesions either in their membrane domain (ERAD-M) or on the ER luminal side (ERAD-L), and each of these pathways is thought to involve different subunits of the complex (*14*). However, as these concepts were developed using mostly artificial model substrates, it is unclear to which extend these pathways degrade endogenous substrates and how these are selected for degradation in the absence of folding lesions. To address these questions, we first defined the *bona fide* substrates of the Hrd1 complex. We found three previously described (Erg3, Erg5, Erg25) (*6, 15*) and two new short-lived proteins (Yjr015w and Ysp2; **Fig. 3B**) that were stabilized in at least five mutants of the Hrd1 complex, and that were also stabilized in cells expressing only a catalytically inactive version of Hrd1 (C399S) (**Fig. 3C**) (*16*). As an example, we further analyzed Erg25 (previously not demonstrated as an Hrd1 target) and found that it interacts with Hrd1 complex members as well as downstream machinery involved in its degradation (**Suppl. Fig S3A**). To test whether endogenous substrates access the ERAD-M or ERAD-L pathway, we analyzed the requirement of their degradation for Der1, which was implicated specifically in degradation of ERAD-L model substrates, such as CPY* (*14*). We did not find striking effects of *DER1* deletion on endogenous short-lived ERAD substrates, suggesting that, under standard growing conditions, ERAD-L is not operating. Consistently, we found no proteins with significantly altered turnover in cells expressing a version of Hrd1 (*Hrd1KRK*) that does not allow domain *in vitro* retrotranslocation of model ERAD-L substrates (data not shown;(*17*)). Instead, each endogenous Hrd1 substrate contains transmembrane domains required for recognition by ERAD-M (**Fig. 3D)**. Also, they each require Yos9, a lectin component of the HRD1 complex, suggesting that their degradation depends on glycosylation (**Fig. 3E)**. This model is supported by the stabilization of Erg3, Erg5, and Ysp2 in *htm1*D cells (**Fig. 3E)**, lacking the a-1,2-specific exo-mannosidase that trims a mannose residue from Man8GlcNac2 (Man8) glycans to form Man7GlcNac2 (Man7), a signal recognized by Yos9. Furthermore, database analysis showed that all endogenous Hrd1 substrates are glycosylated in the luminal portion of the proteins where interactions with Yos9 can occur (**Fig. 3D**; (*18*)). Taken together, these data show that endogenous substrates of the HRD1 complex are degraded by the ERAD-M pathway, likely being recognized through glycosylation.

**Figure 3:**
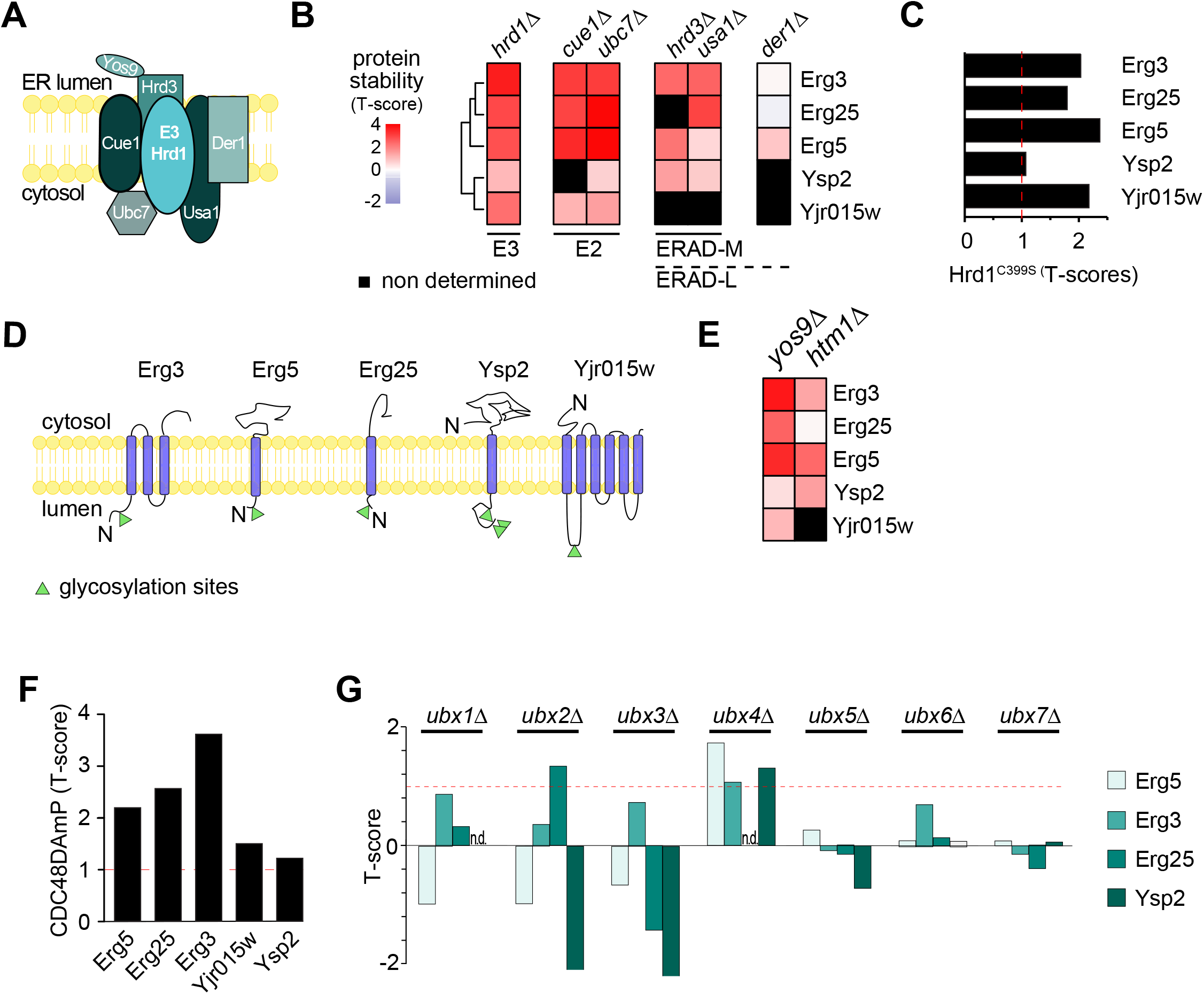
Systematic Prediction of Targets for the HRD1 Branch of ERAD. **(A)** Schematic of the HRD1 branch of ERAD in yeast. **(B)** Hierarchically clustered heat map of the robust T-scores of Hrd1 targets in response to deletion of the member of the HRD complex. **(C)** Stability of Hrd1 targets in cells expressing *Hrd1^C399S^*, a catalytically dead allele of *HRD1*. **(D)** Localization and topology of *bona fide* substrates of the HRD1 complex. Glycosylation sites are indicated as green triangles. **(E)** Heat map clustered as in panel (A) of the robust T-scores of Hrd1 targets in response to deletion of *YOS9* and *HTM1*. **(F)** Stability of HRD1 complex substrates in cells expressing *cdc48DamP*, a hypomorphic allele of *CDC48*. **(G)** Stability of Erg5, Erg3, Erg25 and Ysp2 in cells deleted for individual UBX domain-containing substrate adaptors of Cdc48p.

To further dissect the HRD1-dependent ERAD pathway, we analyzed the requirements for downstream effectors in targeting endogenous substrates for degradation. Remarkably, turnover of Hrd1 targets required the activity of the AAA-ATPase Cdc48 - involved in protein extraction from the membrane (**Fig. 3F**) - and as expected required proteasomal activity (**Suppl. Fig S3B**). Cdc48 is thought to be recruited to substrates by one of the seven Cdc48-binding UBX domain-containing proteins (Ubx1-7). Surprisingly, different Hrd1 targets required different UBX-proteins. Ubx2, is commonly accepted to be the canonical UBX-protein involved in ERAD. However, it was only required for degradation of Erg25, whereas Ubx4 was the predominant factor involved in the degradation of Erg3, Erg5, and Ysp2 (**Fig. 3G)**. Although the mechanistic basis for this difference is unknown, these examples illustrate the power of the T-MAP approach to discover specific degradation pathway requirements.

The T-MAP also provides a powerful approach to determine the biological function of a specific protein degradation pathway. For instance, each of the Hrd1 targets in the T-MAP is involved in sterol metabolism. Erg3, Erg5, and Erg25 are enzymes of ergosterol *de novo* synthesis (*15, 19-21*), *YJR015W* encodes a protein with strong homology to human ABC transporters mediating cellular efflux of sterols (HHPred, **Suppl. Fig S3C**) and Ysp2/Ltc4 belongs to the recently discovered family of StART-domain proteins that localize at membrane contact sites and are responsible for transport of sterols (*22, 23*). These findings expand our understanding of Hrd1-dependent ERAD not only as a master regulator of sterol de novo synthesis, but also its transport in cells.

In addition, we found that key proteins involved in phospholipid (Tgl5, Plb3) and sphingolipid (Tsc10, Orm2, Aur1, Sur1, Csg2, Ipt1) metabolisms are short- or medium-lived. (**Fig. 4A**). Surprisingly, analyses of the T-MAP revealed that these proteins are targeted by different degradation pathways localized in different organelles. Sur1 and Csg2 are proteins in a complex involved in the synthesis of complex sphingolipids (*24*). Both were stabilized when vacuolar degradation or the ESCRT machinery delivering cargoes to vacuole was impaired (**Fig. 4B** and data not shown). In contrast, the regulatory protein Orm2, thought to be localized primarily in the ER (*25*), is targeted by the Tul1 E3 ligase complex to degradation by the proteasome. Tul1 complex shares similarities with the Hrd1 complex but is primarily localized in membranes of the Golgi apparatus and of the endolysosomal system (*26*) (**Fig. 4B, C)**. Finally, degradation by the proteasome of Tsc10, an essential ER and lipid droplet enzyme (*27*) catalyzing the reduction of keto-dihydrosphingosine to dihydrosphingosine (*28*) required Ub-adaptors of ERAD (i.e., stabilized in *ubc7*D, *cue1*D, or *ubc4*D), Cdc48, and Asi1, one of the E3 ligases of the ASI branch of ERAD in yeast (**Fig. 4B, D, E**). In agreement with this, Tsc10 degradation is lower in *asi1*D and *asi3*D and, to a lesser extent, in doa*10*D mutants, than with wildtype cells, when new protein synthesis was blocked with cycloheximide (**Fig. 4F** and **Fig. S4)**. A functional link between the Asi1/3 machinery and sphingolipid metabolism is further supported by genetic interaction data with the strongest interactions reported for *ASI1* and *ASI3* with sphingosine kinase (*LCB5*), sphingosine lyase (*DPL1*) and a component of serine-palmitoyl-CoA transferase (*TSC3*) (*29*) (**Fig. 4G**). These results identify new potential nodes of regulation by degradation control for key enzymes in the sphingolipid metabolism.

**Figure 4:**
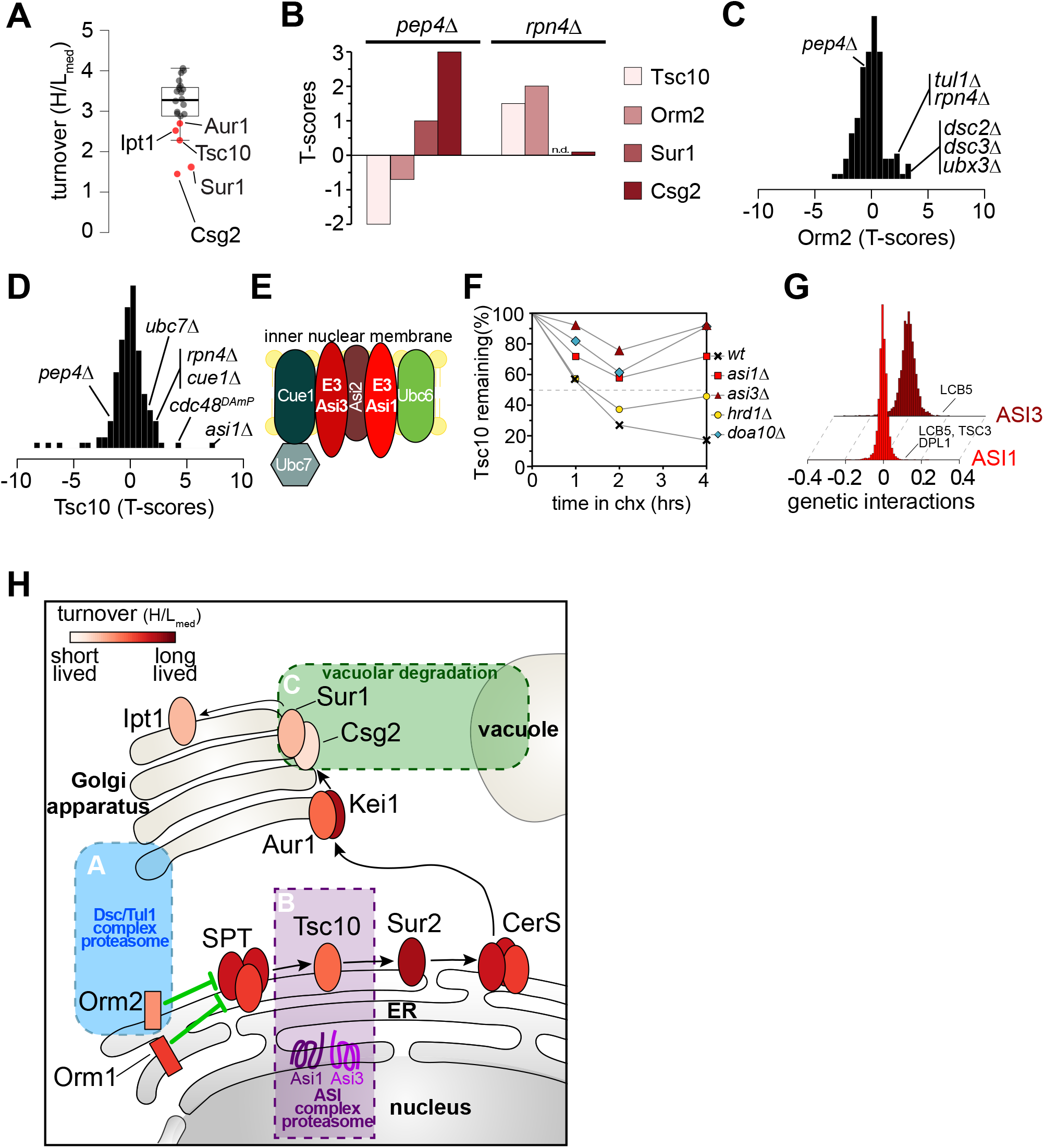
Spatial Control of Degradation for Key Enzymes in the *De Novo* Sphingolipid Biosynthesis Pathway. **(A)** Turnover distribution of enzymes in the *de novo* sphingolipid biosynthesis pathway. **(B)** Stability of Tsc10, Sur1, Csg2, and Orm2 under proteasomal and vacuolar degradation inhibition. **(C)** Stability scores of Orm2 in the mutant present in T-MAP. **(D)** Stability scores of Tsc10 in the mutant present in T-MAP. **(E)** Schematic of the ASI branch of ERAD in yeast. **(F)** Degradation of GFP-TSC10 after inhibition of protein synthesis by cycloheximide in wildtype, Asi1Δ, Asi3Δ, Doa10Δ, and Hrd1Δ. The graph shows the quantification of the western blot shown in Fig. S4. **(G)** ASI1 and ASI3 genetically interact with genes involved in sphingolipids metabolism (Costanzo et al., 2016). **(H)** Model for spatial control of degradation of key enzymes in the *de novo* sphingolipid biosynthesis pathway by the Tul1 complex (blue box), ASI1/3 complex (purple box) and the vacuole (green box). Ellipses represent sphingolipid metabolic enzymes and are color coded according to their half-lives.

In summary, we present a novel analytical platform that combines gene deletions with quantitative measures of protein turnover to deconvolve degradation pathways of *S. cerevisiae*. This particular T-MAP, focusing on protein degradation machinery, provides a rich, new resource for identifying and mechanistically dissecting the ubiquitin-mediated protein degradation system. It reveals endogenous target candidates of nearly all E2 and E3 ligases, as well as the scope of specific pathways, such as ERAD. The analysis of turnover of specific short/medium-lived proteins reveals their degradation pathway and can be utilized to reveal previously unknown nodes of regulation in key cellular metabolisms as underscored by our analysis of sphingolipid metabolism (**Fig. 4H**). Furthermore, the T-MAP approach applied here should be broadly applicable to the study of other cellular processes and systems. For example, extension to other gene sets could reveal target specificities of other components of the proteostasis network (e.g., protein folding or trafficking). Moreover, extension of T-MAP analyses to other organisms, including mammalian systems, provides a powerful new tool for investigating how protein turnover functions in normal physiology and disease. Finally, turnover profiling approach will enable efforts in drug discovery to delineate the therapeutic potential of E3 ligase-modulating agents by allowing the rapid identification of targeted neo-substrates.

## SUPPLEMENTAL INFORMATION

Supplemental Information includes three figures, four tables, and two Treeview sets.

## AUTHOR CONTRIBUTIONS

R.C., T.W.C. designed and supervised the project. R.C. and S.K. carried out and analyzed experiments. R.C., N.M. and T.C.W. performed large-scale analysis and interpretation. R.C. created figures. R.C., N.M, R.V.F., and T.W.C. interpreted the data and wrote the manuscript.

## ACKNOWLEDGMENTS

We would like to thank Drs. Wade Harper and Dan Finley, as well as members of the Farese & Walther laboratory for critical discussion and comments on the manuscript. We thank Drs. Tom Rapoport and Ryan Baldridge for sharing reagents, Xiuling Guo for technical assistance and Gary Howard for editorial help. This work was supported by a grant of the G. Harold and Leila Y. Mathers foundation, NIH GM097194 and the Howard Hughes Medical Institute.

## EXPERIMENTAL PROCEDURES

### Cell Culture and Pulsed SILAC Experiments

Yeast strains were grown in synthetic medium containing 6.7 g/l yeast nitrogen base, 2 g/l dropout mix (US Biological) containing all amino acids except lysine, and 2% glucose. For heavy prelabeling, heavy [^13^C_6_/^15^N_2_] L-lysine (Cambridge Isotope Labs) was added to a final concentration of 30 mg/l. Cells were pre-cultured in 5 ml of medium containing heavy lysine overnight at 30°C and repeated twice. pSILAC experiments were performed by transferring exponentially growing heavy labelled cells, after three washes at 4°C with cold SILAC medium without lysine, to a fresh growth medium containing an excess of light lysine. Cells were harvested at 60 min and immediately prepared for MS analysis. At least two independent biological replicates for each strain were measured.

### Preparation of haploid yeast deletion strains

To minimize compensatory mutations that might accumulate over time in the gene deletion strains, we freshly prepared all mutant strains. Haploid cells harboring the desired deletions were prepared from sporulation of the corresponding BY4743 heterozygous knockout diploid cells (Dharmacon) and selected to be isogenic to parental wild-type BY4742 genetic background (MATα his3Δ1, leu2Δ1, lys2Δ1, ura3Δ1). The deletion and the correct insertion of the kanamycin-resistance cassette were verified by two PCRs using primers described at the Saccharomyces Genome Deletion Project Web page (http://www-sequence.stanford.edu/group/yeast_deletion_project/deletions3.html). Missing haploid knock out strains, even after several rounds of sporulation and selection (28 strains), were directly purchased from Dharmacon and confirmed prior to further analysis.

### Dataset filtering

For each gene deletion turnover profiling dataset, we computed quality metrics including: (1) the number of proteins quantified with at least two ratio counts, (2) the presence of the aminoglycoside phosphotransferase enzyme encoded by the kanamycin cassette conferring resistance to G418, and (3) the absence of the protein encoded by the deleted gene. We excluded, from further analysis, mutant turnover profiling datasets with poor proteome coverage (<=2000 proteins identified).

### Sample preparation

For each mutant, ~25 OD units of cells were harvested by centrifugation. Cells were lysed in 200 μl of buffer containing 50 mM Tris/HCl, pH 9.0, 5% SDS and 100 mM DTT for 30 min at 55°C. Lysates were cleared by centrifugation at 17,000g for 10 min. Supernatants were diluted with buffer UA (8 M urea, 0.1 M Tris/HCl, pH 8.5) to a final concentration of 0.5% SDS. Proteins were digested with the endoproteinase LysC, following the protocol for filter-aided sample preparation (12). Briefly, protein extracts were loaded on a 30k centricon filter unit (Amicon) by centrifugation at 14,000g. Samples were washed twice by addition of 200 μl of buffer UA and alkylated for 20 min in the dark with 5.5 mM iodoacetamide (IAA) in 200 μl of buffer UA. Samples were washed an additional four times by adding 200 μl of buffer UA and centrifugation. 60 μl of buffer UA containing 0.5 mg/ml LysC were added to the filter units and incubated at 37°C overnight. Peptides were recovered by centrifugation into a fresh tube and additional elution with 200 μl of 0.5 M NaCl. Samples were acidified by adding trifluoroacidic acid (TFA) and cleared of precipitates by centrifugation at 17,000g for 5 min. Peptide concentration was measured, and 5 μg of peptides were desalted following the protocol for StageTip purification 13. Samples were eluted with 60 μL of buffer B (80% ACN, 0.1% formic acid in H20) and reduced in a Vacufuge plus (Eppendorf) to a final volume of 3 μL. 2 μL of buffer A (0.1 % formic acid in H20) were added, and the resulting 5 μL were injected into the HPLC.

### Chromatography and Mass Spectrometry

Reversed phase chromatography was performed on a Thermo Easy nLC 1000 system connected to a Q Exactive mass spectrometer (Thermo) through a nano-electrospray ion source. Peptides were separated on 50-cm columns (New Objective) with an inner diameter of 75 μm packed in house with 1.9 μm C18 resin (Dr. Maisch GmbH). Peptides were eluted from 50-cm columns with a linear gradient of acetonitrile from 5–27% in 0.1% formic acid for 240 min at a constant flow rate of 250 nl/min. The column temperature was kept at 40°C by an oven (Sonation GmbH, Germany) with a Peltier element. Peptides eluted from the column were directly electrosprayed into the mass spectrometer. Mass spectra were acquired on the Q Exactive in a data-dependent-mode to automatically switch between full-scan MS and up to 10 data-dependent MS/MS scans. The maximum injection time for full scans was 20 ms with a target value of 3,000,000 at a resolution of 70,000 at m/z=200. The 10 most intense multiple charged ions (z≥2) from the survey scan were selected with an isolation width of 1.4Th and fragment with higher energy collision dissociation (HCD 14) with normalized collision energies of 25. Target values for MS/MS were set to 100,000 with a maximum injection time of 120 ms at a resolution of 17,500 at m/z=200. To avoid repetitive sequencing, the dynamic exclusion of sequenced peptides was set to 45 sec.

### Raw MS Data Analysis

The resulting MS and MS/MS spectra were analyzed using MaxQuant (version 1.5.2.8) and its integrated ANDROMEDA search algorithms. Peak lists were searched against the UNIPROT databases for *S. cerevisiae* with common contaminants added. The search included carbamidomethylation of cysteines as fixed modification, and methionine oxidation and N-terminal acetylation as variable modifications. Maximum allowed mass deviation for MS peaks was set to 6ppm and 20ppm for MS/MS peaks. Maximum missed cleavages were 2. The false discovery rate was determined by searching a reverse database. Maximum false-discovery rates were 0.01 both on peptide and protein levels. Minimum required peptide length was six residues. Proteins with at least two peptides (one of them unique) were considered identified. The “match between runs” option was disabled with a time window of 1 min to match identification between replicates.

### Protein Abundance Estimation

Absolute protein abundances as copies per cell were calculated by applying the “proteomic ruler” plugin in Perseus software using the summed intensities of both heavy and light peptides for each protein groups.

### Data Analysis

We developed custom scripts in R/Bioconductor for data analysis and plotting, which are available on request. Proteins were clustered hierarchically, based on a Pearson correlation of turnover robust score in Cluster, and visualized by TreeView.

### Data and Software Availability

The mass spectrometry proteomics data have been deposited to the ProteomeXchange Consortium via the PRIDE [1] partner repository with the dataset identifier PXD008706 (Username, password: ofy6SXBe).

---

**Suppl. Figure 1:**
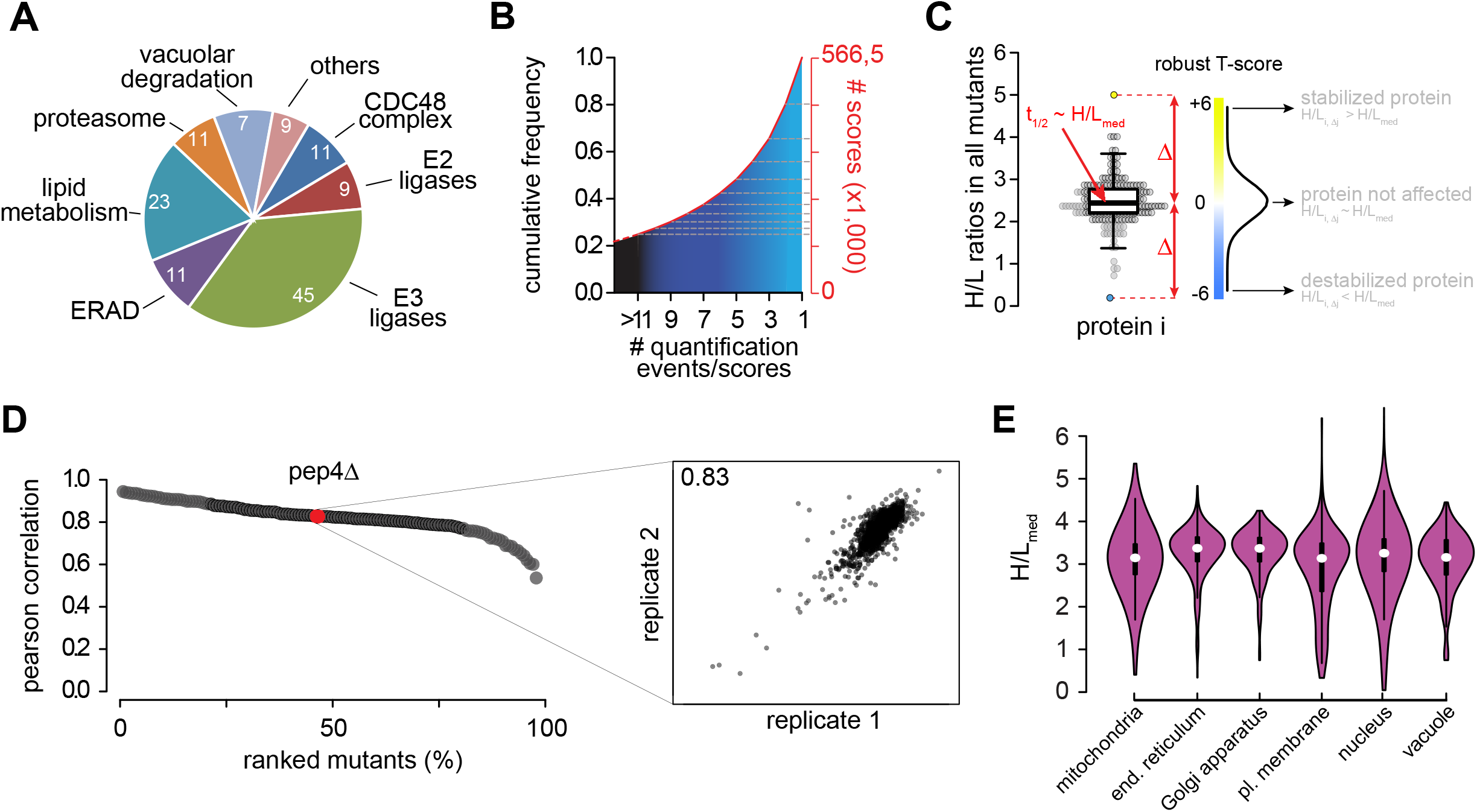
Quality of Turnover Data, Related to Figure 1. **(A)** Pie chart of mutants included in the T-MAP categorized by biological function (described in **Table S1**). **(B)** Cumulative frequency of the number of scores (proteins quantified) in the T-MAP as a function of quantification events/score. **(C)** Scoring function for quantitative turnover analysis identifies destabilizing (H/L_i.Δj_ < H/L_med_), stabilizing (H/L_i.Δj_ > H/L_med_) and neutral (H/L_i.Δj_ ~ H/L_med_) effects of mutant Δ_j_ on protein *i*. **(D)** Reproducibility of turnover profiling between independent biological duplicates (coefficient of correlation) for each mutant in the screen. Inlay shows as an example the reproducibility between biological replicates of *pep4Δ*. **(E)** Distribution of protein turnover (H/L_med_) in different organelles. Abbreviations: end., endoplasmic; pl., plasma.

**Suppl. Figure 2:**
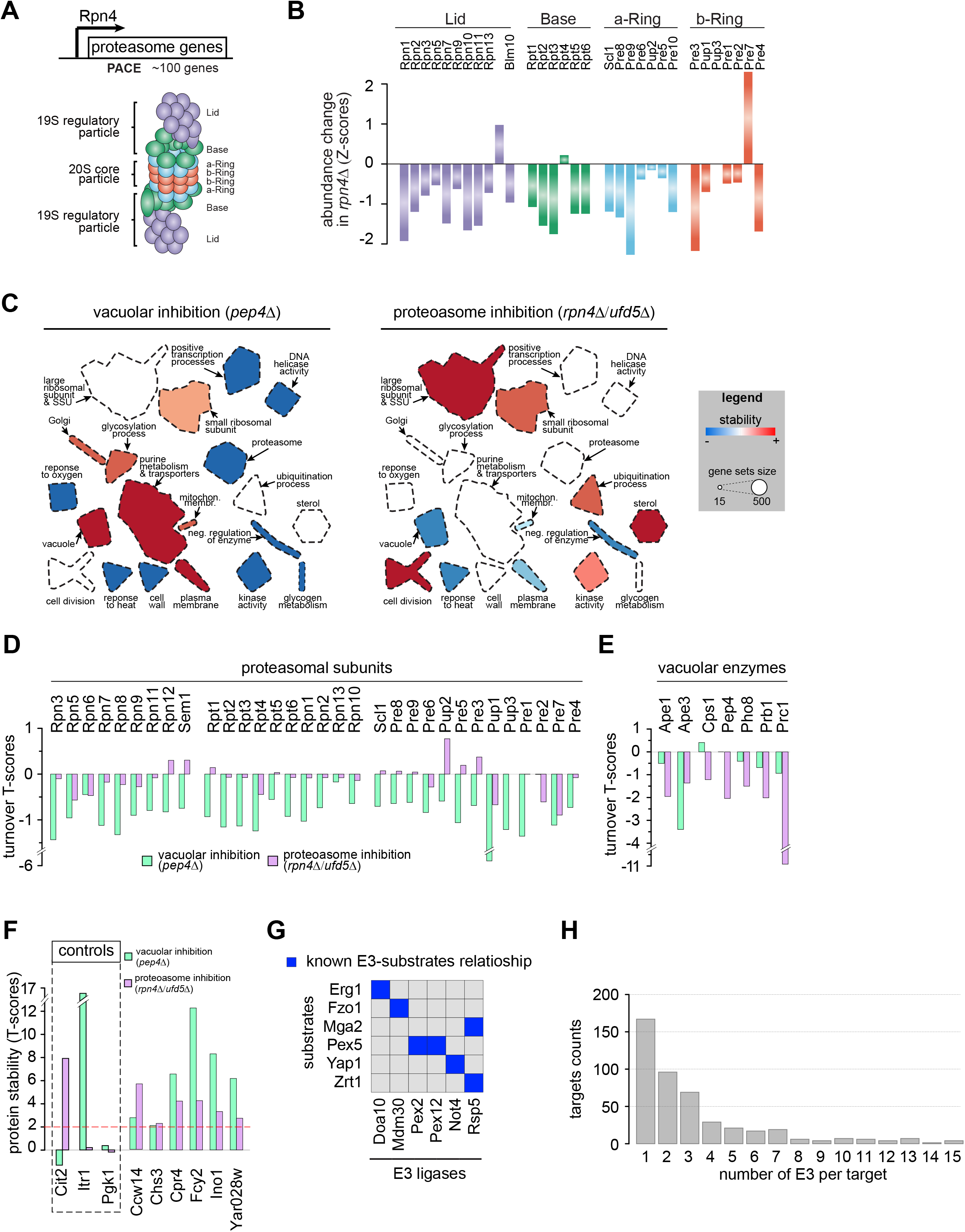
Coordination between Proteasomal and Vacuolar Responses, Related to Figure 2. **(A)** Schematic of the 26S proteasome. The 26S proteasome consists of the catalytic 20S proteasome (a barrel of four stacked rings: two outer rings and two inner rings) and the 19S regulatory particle (RP, also known as PA700). **(B)** Abundance fold-change of proteasome subunit in *rpn4Δ* cells. **(C)** Simplified version of Figure 2A showing separately the effects of vacuolar and proteasomal inhibition on proteome stability. **(D)** Stability (T-scores) of 26S proteasome subunits in *rpn4Δ* and *pep4Δ*. **(E)** Stability (T-scores) of vacuolar enzymes in *rpn4Δ* and *pep4Δ*. **(F)** Six proteins are stabilized by impairing either proteasomal or vacuolar degradation. Itr1 is affected by only vacuolar degradation, Cit2 only by proteasomal degradation, and Pgk1 is an unaffected, long-lived protein. **(G)** Examples of known E3 substrates identified in our T-MAP. Blue indicates known E3 ligase-substrate relationships. **(H)** Distribution of E3 effects on 457 proteins stabilized in at least one E3 ligase mutant strain.

**Suppl. Figure 3:**
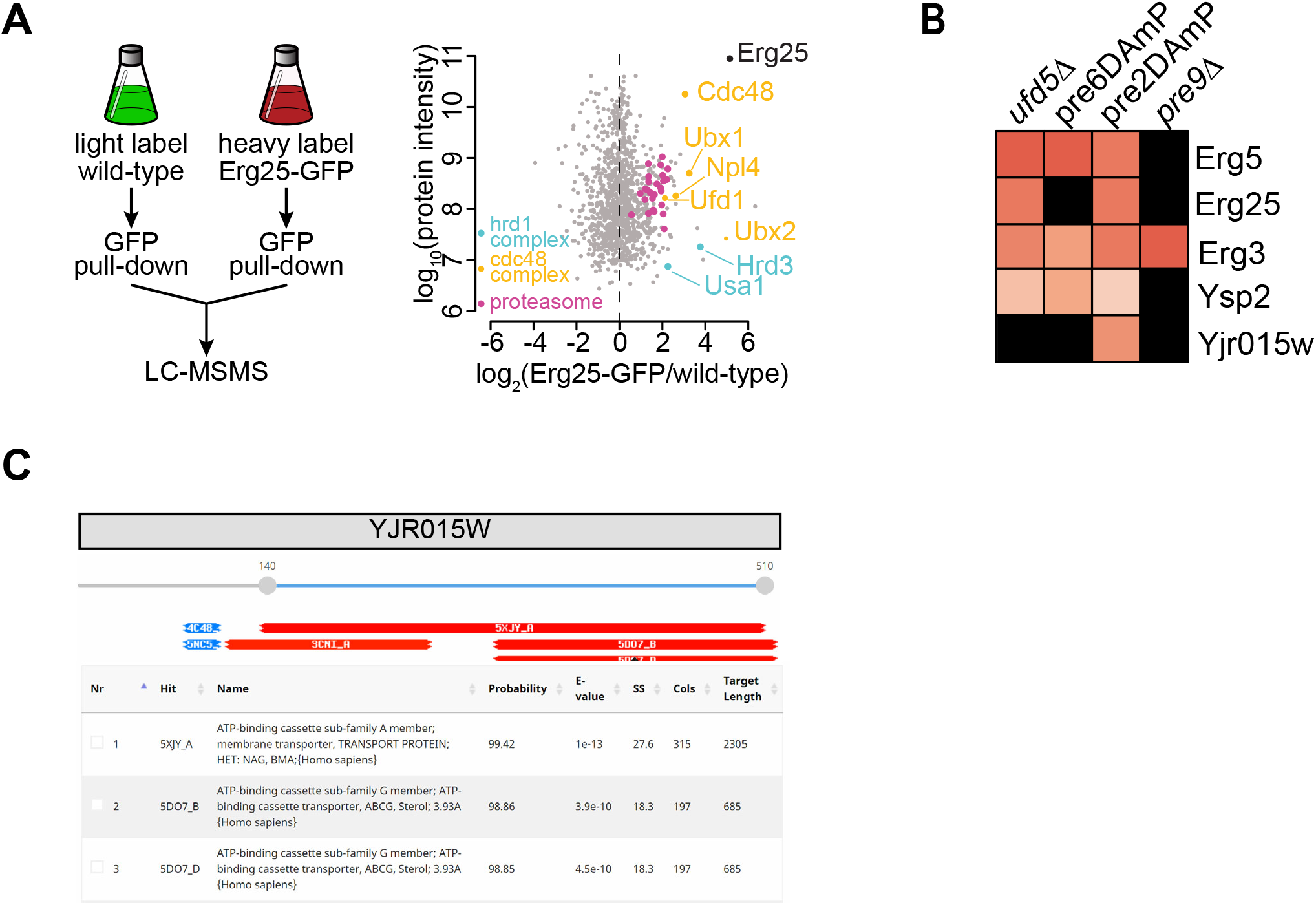
Systematic Prediction of Targets for the HRD1 Branch of ERAD, Related to Figure 3. **(A)** SILAC-based pull down of Erg25 fused to GFP. (left) Erg25 physically interacts with the HRD1 complex, the CDC48 complex, and the proteasome. Experimental design of affinity purification and MS analysis of “heavy”-labeled cells expressing GFP-tagged Erg25 and untagged control cells. (right) Intensities are plotted against normalized heavy/light SILAC ratios **(B)** Stability of Hrd1 targets in cells with impaired proteasomal activity deleted for either rpn4 or pre9 or expressing the DAMP allele of the essential genes PRE2 and PRE6. **(C)** HHPred alignment of YJR015W.

**Suppl. Figure 4:**
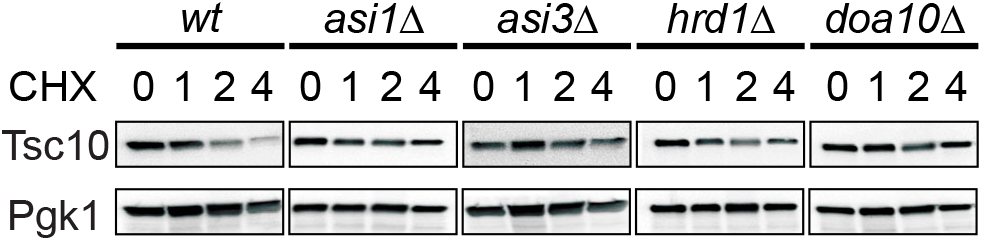
Spatial Control of Degradation for Key Enzymes in the *De Novo* Sphingolipid Biosynthesis Pathway, Related to Figure 4. Degradation of GFP-TSC10 after inhibition of protein synthesis by cycloheximide in wildtype, Asi1Δ, Asi3Δ, Doa10Δ, and Hrd1Δ. Cell extracts were analyzed by SDS–polyacrylamide gel electrophoresis and blotting.

